# Targeted degradation of α-synuclein prevents PFF-induced aggregation

**DOI:** 10.1101/2025.04.15.648905

**Authors:** Bill Carton, Géraldine Gelders, Gajanan Sathe, Nur M. Kocaturk, Sascha Roth, Thomas J. Macartney, Joke Van Elsen, Louis De Muynck, Arjan Buist, Diederik Moechars, Gopal P. Sapkota

**Affiliations:** MRC Protein Phosphorylation and Ubiquitylation Unit, School of Life Sciences, University of Dundee, Sir James Black Centre, Dow Street, Dundee, DD1 5EH, United Kingdom; Neuroscience Discovery, Johnson & Johnson, Turnhoutseweg 30, 2340 Beerse, Belgium

## Abstract

Accumulation of misfolded α-synuclein protein in intracellular inclusion bodies of dopaminergic neurons underlies the pathogenesis of Synucleinopathies, which include Parkinson’s Disease (PD), Dementia with Lewy Bodies (DLB) and Multiple System Atrophy (MSA). Therefore, clearance of misfolded α-synuclein from dopaminergic neurons could in principle offer a therapeutic window for Synucleinopathies, which currently remain untreatable. In this study, we employ the Affinity-directed PROtein Missile (AdPROM) system consisting of the substrate receptor of the CUL2-E3 ligase complex VHL and a nanobody selectively recognising the human α-synuclein protein and demonstrate targeted degradation of endogenous α-synuclein from human cell lines with remarkable selectivity. We further demonstrate that targeted degradation of α-synuclein prevents the pre-formed fibril (PFF)-induced aggregation of α-synuclein in primary neurons derived from rats expressing human α-synuclein. This approach represents the first demonstration of nanobody-guided proteasomal degradation of all clinically relevant α-synuclein variants, highlighting its potential as a therapeutic strategy against Synucleinopathies.

## Introduction

Intracellular inclusions containing the small 140-amino acid protein α-synuclein are found in dopaminergic neurons in the substantia nigra pars compacta of patients with a number of different progressive neurodegenerative disorders, such as Parkinson’s Disease (PD), Dementia with Lewy Bodies (DLB) and Multiple System Atrophy (MSA), known collectively as synucleinopathies^1–4^. Effective treatments to cure synucleinopathies remain elusive, and with the increase in life expectancy globally, the need for them has become critical. Amongst synucleinopathies, PD is the most common, and, according to Parkinson’s Europe, 10 million people worldwide currently live with PD. PD is a movement disorder and the main symptoms include bradykinesia, resting tremor and postural instability as well as reported cognitive symptoms such as depression and dementia^5^. The underlying molecular causes of synucleinopathies remain unknown, although the Lewy bodies are characteristic hallmark of the diseases and may contain potential therapeutic targets. The Lewy bodies consist of a vast number of different proteins but the primary constituent is α-synuclein, which is a protein encoded by the *SNCA* gene^6^. Strong causal links between α-synuclein and dominantly inherited forms of PD and DLB also exist, because several missense mutations in the *SNCA* gene (including A30P, E46K, H50Q, G51D, A53E and A53T) as well as multiplication of the *SNCA* gene are known to be associated with pathology^7–12^. Although *SNCA* mutations account for a low proportion of familial PD cases, misfolded and aggregated forms of the α-synuclein protein are found in about 95% of all PD brains^13–16^. Therefore, removing or reducing the levels of toxic forms of α-synuclein from patient neurons potentially offers a therapeutic opportunity for synucleinopathies.

The precise physiological functions of α-synuclein and its contribution to the pathological progression of synucleinopathies remain to be elucidated. The α-synuclein protein abundance is enriched at presynaptic terminals^17^ and research suggests that it plays a role in SNARE complex formation allowing for synaptic vesicle release^18^. As a member of the intrinsically disordered protein family, α-synuclein itself has no tertiary structure and its conformation can be affected by a number of different factors, such as the presence of interactors or lipid membranes^19^. Within the Lewy bodies, the misfolded α-synuclein is heavily phosphorylated at Ser129, and this phosphorylation is often used as a marker to confirm the presence of misfolded/aggregated structures, as only trace levels of the phosphorylated α-synuclein protein are evident in the brains of healthy individuals compared to patients with PD^20,21^. However, it is unclear whether this phosphorylation is involved in the aggregation process or pathogenesis of PD. Whether the toxicity elicited on affected neurons is due to α-synuclein aggregates themselves, via a toxic gain of function, or due to disruption of α-synuclein function from its endogenous site of action, thereby eliciting a loss of function, is also yet to be determined. Nevertheless, the strong evidence linking aggregated α-synuclein to the pathogenesis and progression of PD^1–4^, along with *SNCA* mutations and multiplications leading to α-synuclein aggregation and PD pathology^7–12^ make α-synuclein a promising target for therapeutic intervention. Indeed, several studies have reported that lowering of the α-synuclein protein abundance through targeting the *SNCA* gene and transcript has beneficial effects in PD disease models^22–24^. In the absence of a known biological activity and disordered nature of protein conformation, α-synuclein has been considered an intractable target through conventional small-molecule drug discovery^25^. Therefore, new, and innovative approaches to target α-synuclein are required for both the advancement of research and the development of novel therapeutic approaches.

Targeted protein degradation (TPD) has emerged as an effective modality for targeting the elimination of proteins of interest (POI)^26,27^. Indeed, some small molecule degraders, such as Proteolysis targeting chimeras (PROTACs) and molecular glues, are already in pre-clinical or clinical use for different indications^27^. There are many TPD approaches that harness cellular proteolytic pathways^27,28^. One such modality is the proteolytic affinity-directed protein missile (AdPROM) system, which utilises the ubiquitin proteasome system to target the degradation of proteins in cells^28–31^. Briefly, the proteolytic AdPROM comprises an E3 ligase or an E3 ligase substrate receptor, such as the von Hippel–Lindau (VHL) protein, connected to a selective polypeptide binder, such as a nanobody, of the target protein. When delivered to a cell, the AdPROM system enables the proximity between the target protein and the E3 ligase facilitating the ubiquitylation and degradation of the target protein. The AdPROM system has been employed to efficiently and selectively degrade many intracellular target proteins in different human cell lines^28,30,31^. In this study we explore targeted degradation of α-synuclein through the proteolytic AdPROM system.

## Results

### VHL-aGFP16 and VHL-NbSYN87 AdPROMs degrade GFP-tagged α-synuclein overexpressed in U2OS osteosarcoma cells

In order to evaluate whether α-synuclein could be targeted for degradation by the proteolytic AdPROM, we first generated Flp-IN T-Rex U2OS osteosarcoma cells in which a single copy of the GFP-tagged α-synuclein, either the wild type (WT) or the pathogenic A53T mutant, was integrated into a specific genomic locus containing an upstream tetracycline-inducible promoter. Treatment of these cells with doxycycline induced an increase in GFP-tagged α-synuclein expression over time but in both cases some protein expression was detected even in the absence of doxycycline. Substantially more GFP-tagged mutant A53T α-synuclein was detected relative to WT α-synuclein (Fig. S1A). Using the previously validated VHL-aGFP16 AdPROM^28,30,31^ as well as the newly generated VHL-NbSYN2 and VHL-NbSYN87 AdPROMs that incorporate nanobodies reported to interact with α-synuclein^32^, we sought to demonstrate whether GFP-tagged WT and A53T α-synuclein in these cells could be targeted for degradation (Fig. 1A-C). The levels of both GFP-tagged WT and A53T α-synuclein in cells transduced with VHL-aGFP16 AdPROM expressing retroviral vectors were substantially lower than in untransduced cells or those transduced with VHL or aGFP16 alone controls (Fig. 1D&E). Excitingly, the levels of both GFP-tαgged α-synuclein forms in cells transduced with VHL-NbSYN87 AdPROM were substantially decreased compared to untransduced cells or cells transduced with NbSYN87 and VHL controls, suggesting potential targeted degradation of GFP-tagged α-synuclein (Fig. 1D&E). The VHL-NbSYN2 AdPROM did however not change the levels of GFP-tagged WT and A53T α-synuclein relative to untransduced cells or cells transduced with NbSYN2 and VHL controls (Fig. 1D&E). Compared to VHL-aGFP16 and VHL-NbSYN87, the levels of VHL-NbSYN2 were substantially lower, although all cells were cultured in selection medium containing puromycin to select for the transduction of the retroviral vectors encoding these AdPROMs. The ability of VHL-NbSYN87 to efficiently degrade GFP-tagged α-synuclein, although exciting, could be mediated through ubiquitylation of lysine residues on the GFP tag rather than on the α-synuclein itself. Therefore, we sought to test the effect of VHL-NbSYN87 to degrade the untagged WT and pathogenic mutant forms of α-synuclein (Fig. 2A&B).

**Figure 1:**
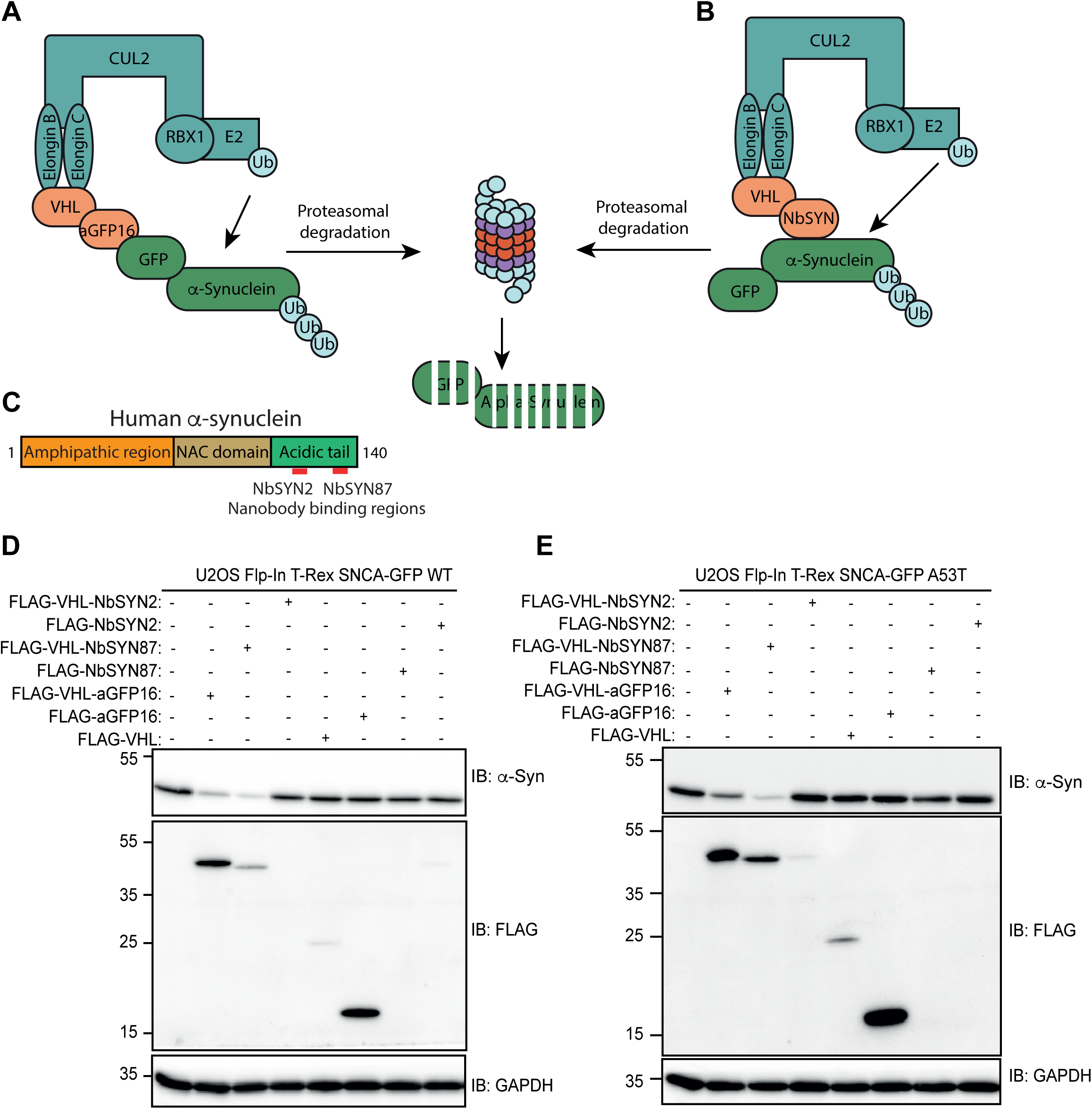
Targeted degradation of GFP-α-synuclein through anti-GFP16 and anti-α-synuclein nanobody AdPROMs. **A)** Schematic describing the targeted degradation of GFP-tagged α-synuclein by VHL-aGFP16 AdPROM, in which the aGFP nanobody recruits GFP-tagged α-synuclein to the Cullin 2 RING E3 ligase complex via VHL for its polyubiquitination and subsequent degradation through the proteasome. **B)** As in A, except the schematic shows targeted degradation of α-synuclein through VHL-NbSYN AdPROM, in which a nanobody targeting the α-synuclein (NbSYN) replaces the aGFP16 nanobody. **C)** Domain architecture of human α-synuclein, with the approximate recognition sites for the NbSYN2 and NbSYN87 nanobodies indicated. NAC – Non-Amyloid Component. **D&E)** U2OS Flp-In T-Rex cells expressing GFP-tagged WT **(D)** or A53T **(E)** α-synuclein were retrovirally transduced to express either the VHL-aGFP16, VHL-NbSYN2 or VHL-NbSYN87 AdPROMs or the indicated controls. 20 μg of extract protein was resolved by SDS-PAGE and transferred onto nitrocellulose membranes, which were subjected to immunoblotting with the indicated antibodies. All blots representative of at least n=3.

**Figure 2:**
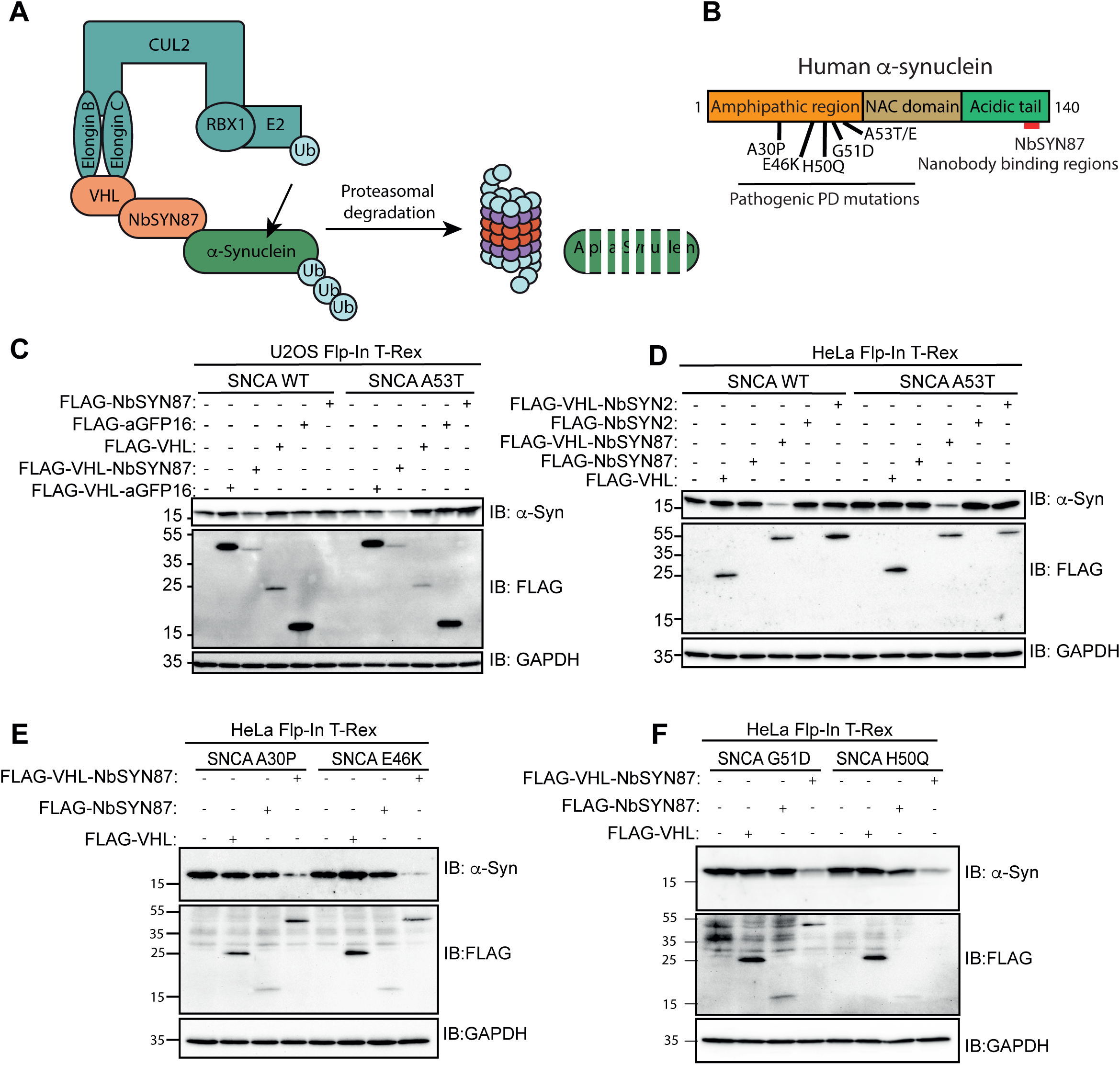
Targeted degradation of untagged α-synuclein through the VHL-NbSYN87 AdPROM. **A)** Schematic depicting the use of VHL-NbSYN87 AdPROM to induce polyubiquitination of untagged α-synuclein through the proteasome. **B)** Domain architecture of human α-synuclein, with the reported familial PD pathogenic mutations indicated. Approximate recognition site for the NbSYN87 nanobody is also indicated. **C)** U2OS Flp-In T-Rex cells expressing either WT or A53T mutant α-synuclein were retrovirally transduced to express either VHL-NbSYN87 or VHL-aGFP16 alongside the indicated controls. 20 μg of extract protein was resolved by SDS-PAGE and transferred onto nitrocellulose membranes, which were subjected to immunoblotting with the indicated antibodies. Blots representative of at least n=3. **D)** As in (C), except HeLa Flp-In T-Rex cells expressing either human WT or A53T α-synuclein were employed. **E)** HeLa Flp-In T-Rex cells expressing either A30P or E46K mutants of human α-synuclein were retrovirally transduced to express VHL-NbSYN87, VHL, or NbSYN87. 20 μg of extract protein was resolved by SDS-PAGE, transferred onto nitrocellulose membranes, which were subjected to immunoblotting with the indicated antibodies. Blots representative of at least n=3. **F)** As in (E) except Flp-In T-Rex cells expressing G51D and H50Q mutants of human α-synuclein were used.

### VHL-NbSYN87 AdPROM degrades wild type and pathogenic PD mutants of α-synuclein overexpressed in HeLa cells

We generated Flp-In T-Rex cells stably integrated with untagged WT and mutant A53T α-synuclein in both U2OS (Fig. 2C) and HeLa cells (Fig. 2D). Transduction with the VHL-aGFP16 AdPROM did not show reduced levels of untagged WT nor A53T α-synuclein in both cell lines in comparison to untransduced cells, or those transduced with VHL or aGFP16 alone controls (Fig. 2C&D). These findings suggest that the aGFP16 nanobody, as expected, does not target untagged α-synuclein. Interestingly, both U2OS and HeLa cells transduced with VHL-NbSYN87 AdPROM showed a substantial decrease in levels of both WT and A53T α-synuclein when compared to untransduced cells or those transduced with VHL, or NbSYN87 controls (Fig. 2C&D). In contrast, cells transduced with VHL-NbSYN2 did not yield a reduction in either WT and A53T α-synuclein levels in comparison to untransduced cells or those transduced with NbSYN2 or VHL alone controls (Fig. 2C&D). These results indicate that the VHL-NbSYN87 is capable of degrading untagged WT and A53T α-synuclein overexpressed in both U2OS and HeLa cell lines. Next, we sought to study the ability of the VHL-NbSYN87 AdPROM to degrade four other mutants of the α-synuclein protein reported in familial cases of PD (Fig. 2B). HeLa Flp-In T-Rex cells expressing each of the four α-synuclein mutants, namely A30P, E46K, G51D and H50Q (Fig. S1B), displayed a substantial reduction in the levels of all of the α-synuclein mutants upon transduction with VHL-NbSYN87 AdPROM compared to untransduced cells or those transduced with NbSYN87 and VHL alone controls (Fig. 2E&F). Collectively, these observations demonstrate that the VHL-NbSYN87 AdPROM enables the degradation of untagged α-synuclein and all of the clinically relevant mutated forms of the protein tested above.

### VHL-NbSYN87 AdPROM degrades endogenous α-synuclein

Having established that VHL-NbSYN87 AdPROM is able to degrade exogenously expressed α-synuclein, we sought to assess whether it could target the degradation of endogenous α-synuclein in distinct clonal cell lines. To this end, we first turned to routinely used laboratory human cell lines, primarily derived from cancers, to monitor the levels of endogenous α-synuclein expression (Fig. 3A, 3B & S2). From this screen, we found SK-MEL13, SK-MEL-23 and G-361 melanoma cells, H23 epithelial-like cells, H460 non-small cell lung cancer cells, SH-SY5Y neuroblastoma cells and HeLa cervical cancer cells to express robust levels of α-synuclein that were easily detectable by immunoblotting (Fig. 3A&B and Fig. S3). The 24 other cells lines we tested only expressed low levels of α-synuclein or levels that were not detectable by immunoblotting (Fig. 3A&B and Fig. S2A&B). When SK-MEL-13 and G-361 melanoma cells were transduced with the VHL-NbSYN87 AdPROM, there was almost complete loss in levels of endogenous α-synuclein compared to untransduced cells or those transduced with VHL-aGFP16, VHL, aGFP16 or NbSYN87 controls (Fig. 3C-E). Compared to control cells, the expression of VHL-NbSYN87 in SK-MEL-13 cells did not cause any change in the levels of *SNCA* transcripts (Fig. S3). These results suggest that the VHL-NbSYN87 AdPROM targets the degradation of physiological α-synuclein protein from cells. In order to determine whether the loss in levels of endogenous α-synuclein by VHL-NbSYN87 was indeed mediated by the CUL2 E3 ligase machinery, for which VHL is the substrate receptor^33^, we treated G-361 cells with the pan-Cullin neddylation inhibitor MLN4924^34^ for 24 hours prior to cell lysis. As expected, treatment of G-361 cells with MLN4924 resulted in the inhibition of CUL2 neddylation and a concurrent stabilization of its endogenous target, HIF1α (Fig. 3F). Under these conditions, the loss in levels of endogenous α-synuclein caused by the VHL-NbSYN87 AdPROM was partially rescued by MLN4924 (Fig. 3F), suggesting the requirement of CUL2 E3 ligase activation. Importantly, the expression of VHL-NbSYN87 had no impact on the levels of endogenous HIF1α relative to controls, suggesting the AdPROM system does not interfere with the endogenous CUL2 E3 ubiquitin ligase machinery (Fig. 3F).

**Figure 3:**
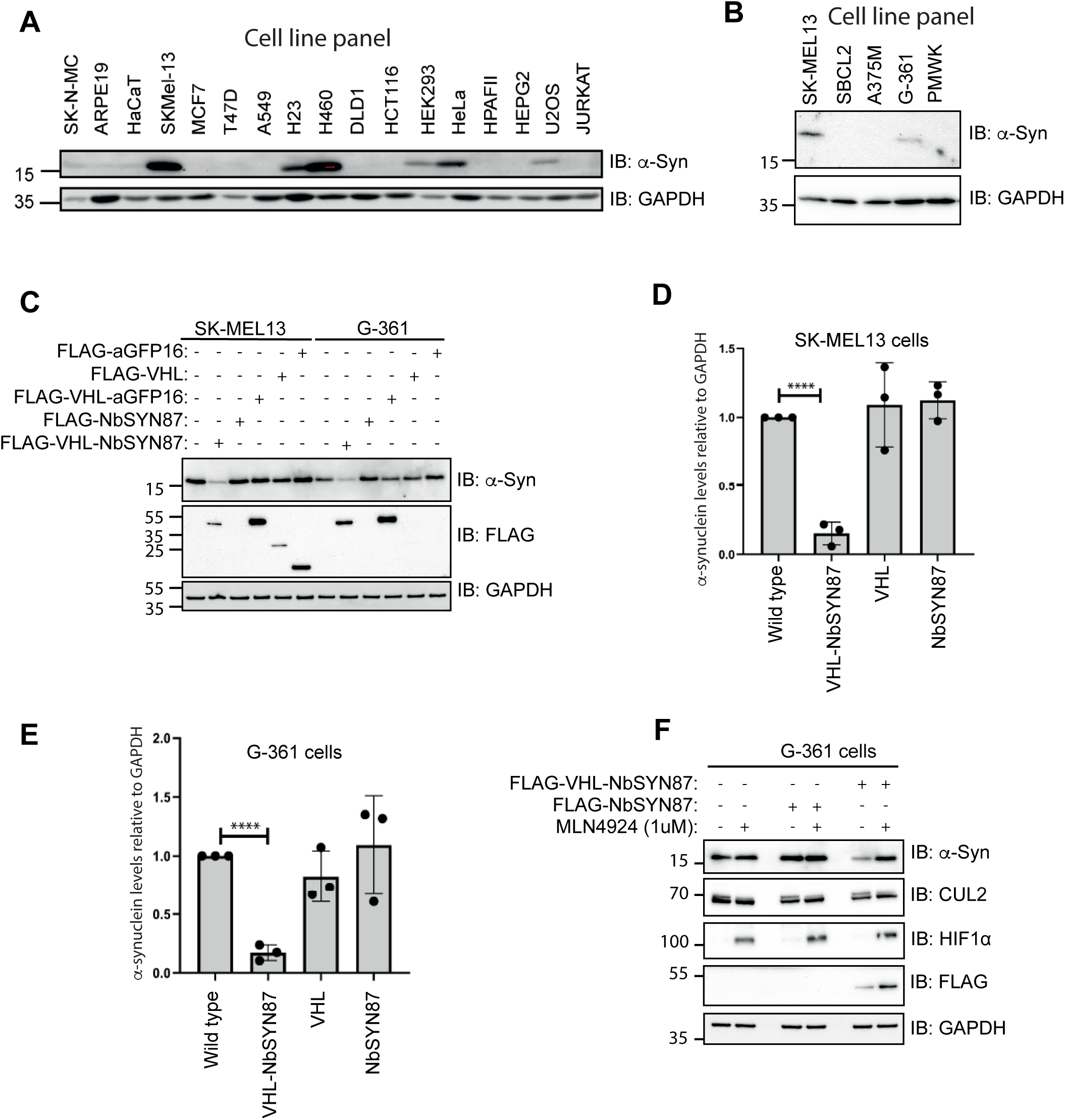
Targeted degradation of endogenous α-synuclein in melanoma cells. **A)** Extracts (20 µg protein) from 17 different human cell lines were resolved by SDS-PAGE and transferred to nitrocellulose membranes for immunoblotting with the indicated antibodies. **B)** Extracts (20 µg protein) from a panel of five different melanoma cells were resolved by SDS-PAGE and transferred to nitrocellulose membranes for immunoblotting with the indicated antibodies. **C)** SK-MEL13 and G-361 cells were retrovirally transduced to express either VHL-NbSYN87 or VHL-aGFP16 as well as the indicated controls. 20 μg of extract protein was resolved by SDS-PAGE and transferred onto nitrocellulose membranes, which were subjected to immunoblotting with the indicated antibodies. Blots representative of at least n=3. **D&E)** Quantitation of relative α-synuclein levels from (C) normalised to GAPDH control ±SD of n=3 independent experiments for SK-MEL-13 **(D)** and G-361 **(E)** cells. Statistical analysis was carried out using a Students T-test. **** = P < 0.0001. **F)** G-361 cells expressing either VHL-NbSYN87 or NbSYN87 alone were treated with 1 μM of the neddylation inhibitor MLN4924 for 24 hours prior to lysis. 20 μg of extract protein was resolved by SDS-PAGE and transferred onto nitrocellulose membranes, which were subjected to immunoblotting with the indicated antibodies. Blots representative of at least n=3.

### Comparing targeted degradation of α-synuclein through VHL-NbSYN87 against genetic silencing approaches

To compare the levels of degradation achieved by the VHL-NbSYN87 AdPROM to an siRNA-mediated depletion of α-synuclein, SK-MEL-13 cells were either transduced with the VHL-NbSYN87 AdPROM or transiently transfected with siRNAs targeting *SNCA* transcripts (Fig. 4A&B). As before, the VHL-NbSYN87 AdPROM resulted in a significant reduction in levels of α-synuclein by ∼95% compared to untransduced control cells (Fig. 4A&B). In comparison, siRNAs targeting *SNCA* transcripts led to a reduction of α-synuclein protein levels by ∼50% compared to control non-targeting siRNAs (Fig. 4A&B). These results imply that the VHL-NbSYN87 AdPROM causes a greater reduction in α-synuclein protein levels in SK-MEL-13 cells compared to targeting *SNCA* transcripts with siRNAs. We also assessed the VHL-NbSYN87-mediated degradation of α-synuclein relative to the genetic ablation of the *SNCA* gene by CRISPR/Cas9 (Fig. 4C-D & Fig. S4). The reduction in α-synuclein protein levels caused by the VHL-NbSYN87 AdPROM was comparable to the complete *SNCA* knockout (KO) cells (Figure 4C-D). The *SNCA*-KO SK-MEL-13 clones allowed us to validate the complete degradation of α-synuclein caused by the VHL-NbSYN87 AdPROM by immunostaining with anti-α-synuclein antibody. This led to the absence of signal in *SNCA*-KO and VHL-NbSYN87 transduced cells compared to a robust signal observed for the WT SK-MEL-13 cells (Fig. 4E).

**Figure 4:**
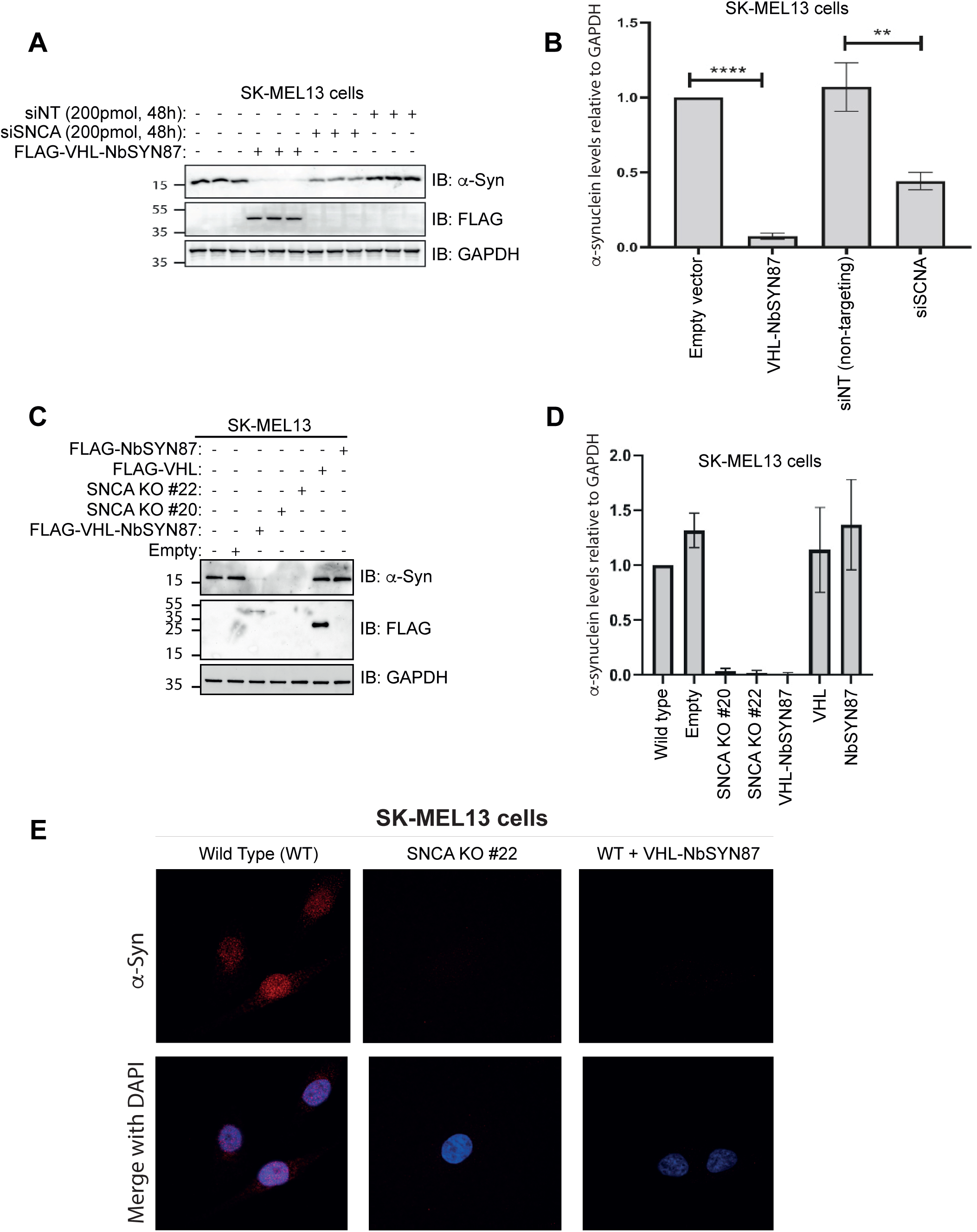
Depletion of α-synuclein in SK-MEL-13 melanoma cells via the VHL-NbSYN87 AdPROM, siRNAs or CRISPR/cas9 genome editing technologies. **A)** SK-MEL-13 cells were retrovirally transduced with empty viral vector or VHL-NbSYN87 AdPROM before lysis. In parallel, SK-MEL-13 WT cells were treated with 20 pmol *SNCA*-targeting siRNA (siSNCA) or with non-targeting siRNA control for 48 hours prior to lysis. Cell extracts (20 µg protein) were resolved by SDS-PAGE and transferred to nitrocellulose membranes before immunoblotted with the indicated antibodies. Blots representative of n=3 independent experiments. **B)** Relative quantification of α-synuclein from (A) normalised to GAPDH controls. Error bars show ±SD of n=3 independent experiments. Statistical analysis was carried out using Students T-test. ****, P-value < 0.0001. **, P-value < 0.01. **C)** SK-MEL-13 cells transduced with the VHL-NbSYN87 AdPROM or controls were lysed alongside two isolates of *SNCA*-KO SK-MEL-13 cells that were generated using a CRISPR/CAS9 strategy. 20 μg of extract protein was resolved by SDS-PAGE and transferred onto nitrocellulose membranes, which were subjected to immunoblotting with the indicated antibodies. Blots representative of n=3 independent experiments. **D)** Quantification of α-synuclein levels from (A) normalised to GAPDH loading control ± SD of n=3 independent experiments. **E)** Representative images (n=3 independent experiments) from immunofluorescence staining of WT SK-MEL-13 cells transduced with empty vector or the VHL-NbSYN87 AdPROM and *SNCA*-KO clone #22 with an anti-α-synuclein antibody (red channel). DNA is stained with DAPI (blue channel).

### Targeted degradation of α-synuclein through VHL-NbSYN87 is highly specific

To determine the specificity of VHL-NbSYN87-mediated degradation of α-synuclein, we assessed the global quantitative proteomic changes upon transduction of the VHL-NbSYN87 AdPROM in SK-MEL-13 cells compared to WT cells transduced with control empty vector.

Immunoblotting of SK-MEL-13 cell extracts, undertaken prior to an unbiased total quantitative proteomics using tandem mass tagging (TMT) approach, verified that the VHL-NbSYN87 AdPROM resulted in almost complete degradation of α-synuclein compared to WT controls, with *SNCA*-KO extracts used as complete α-synuclein depletion control (Fig. 5A). Total quantitative proteomic analysis identified a total of 6906 unique peptides. When changes in abundance between WT SK-MEL-13 cells were compared to those transduced with VHL-NbSYN87 AdPROM, only peptides from two proteins, namely VHL and α-synuclein, showed significant differences. α-synuclein was the only protein whose abundance was found to be significantly downregulated in cells transduced with the VHL-NbSYN87 AdPROM, while VHL was found to be significantly upregulated, which is to be expected due to overexpression of the VHL-NbSYN87 AdPROM system in these cells (Fig. 5B). This highlights the remarkable and exquisite specificity of the degradation achieved with the VHL-NbSYN87 AdPROM. From this data, it also appears that over the duration of α-synuclein degradation by the VHL-NbSYN87 AdPROM, there did not appear to be any other significant proteomic changes in SK-MEL-13 cells as a consequence of α-synuclein degradation. Of note, protein levels of the other two members of the synuclein family, β- and γ-synuclein, were unchanged across samples further indicating the specificity of the VHL-NbSYN87 AdPROM.

**Figure 5:**
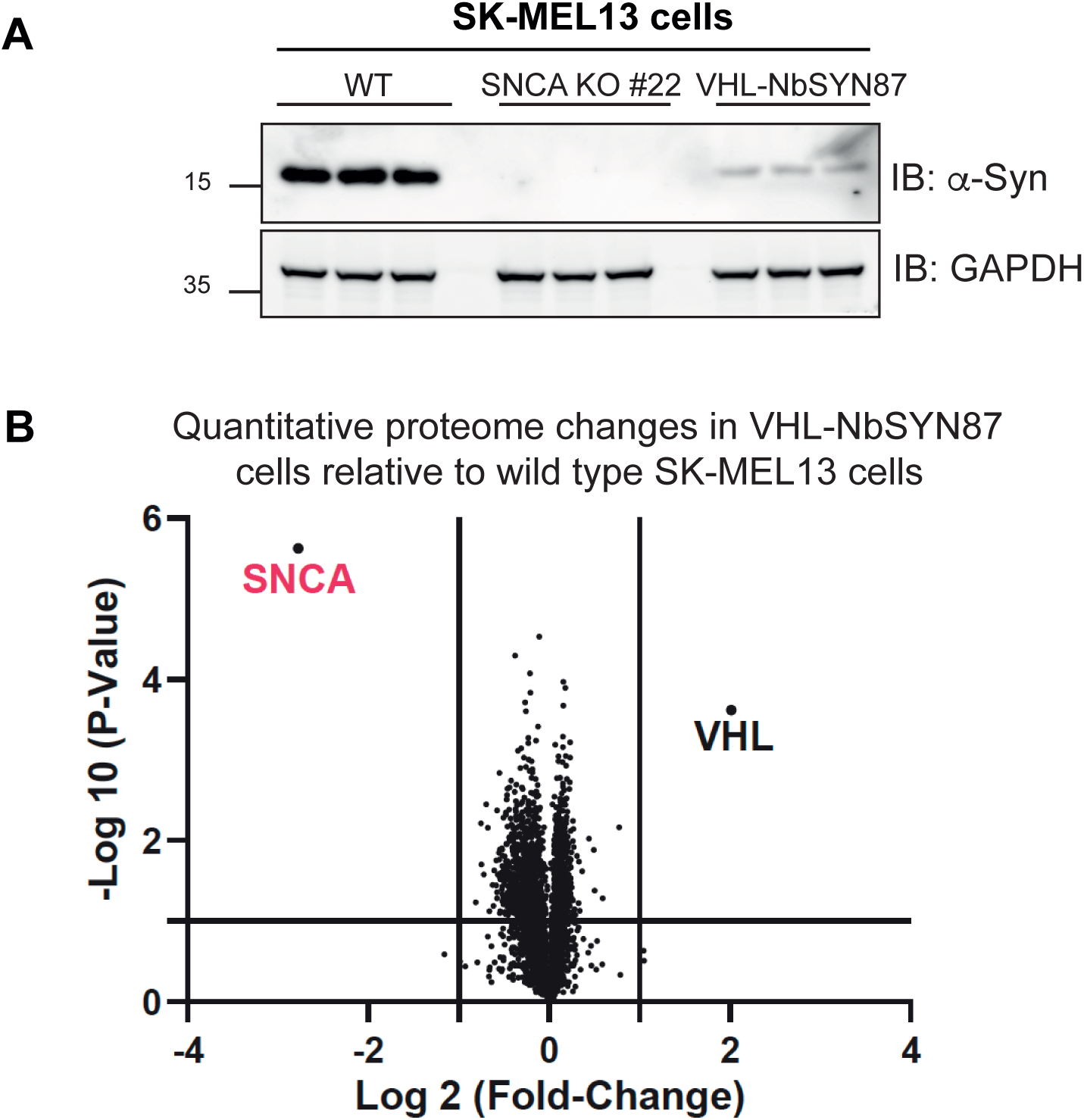
AdPROM-mediated degradation of α-synuclein is highly specific. **A)** WT SK-MEL-13, *SNCA^-/-^* and FLAG-VHL-NbSYN87 transduced WT SK-MEL-13 cells were lysed and cell extracts (20 µg protein) were resolved by SDS-PAGE and transferred to nitrocellulose membranes for immunoblotting with the indicated antibodies. Each of the 3 replicates are included in the blot. **B)** The identical extract samples from (A) were subjected to total quantitative proteomic analysis using TMT. The Volcano plot shows total quantitative proteomic changes evident in extracts between WT SK-MEL-13 cells over those transduced with the VHL-NbSYN87 AdPROM. Horizontal line indicates threshold of statistical significance of P-value = 0.05. Two vertical lines indicate a negative (left) and positive (right) 2-fold changes in proteins.

### The VHL-NbSYN87 AdPROM degrades human but not mouse α-synuclein

We also evaluated the ability of the VHL-NbSYN87 to degrade endogenous α-synuclein in human SH-SY5Y neuroblastoma cells. Cells transduced with VHL-NbSYN87 resulted in a robust and significant reduction in levels of α-synuclein compared to those transduced with empty vector, VHL, or NbSYN87 controls alone (Fig. 6A&B). Together with the previous observations across many different cell lines, it is evident that the VHL-NbSYN87 AdPROM is capable of degrading human α-synuclein. Human α-synuclein shares approximately 97% sequence identity with the mouse protein (Fig. 6C). Although NbSYN87 nanobody was generated using the human α-synuclein as the antigen, we tested the ability of VHL-NbSYN87 to degrade mouse α-synuclein. When U2OS Flp-In T-Rex cells expressing either mouse or human α-synuclein were transduced with the VHL-NbSYN87 AdPROM, no degradation of the mouse α-synuclein but near complete degradation of human α-synuclein was observed compared to cells transduced with empty vector, VHL, or NbSYN87 alone controls (Fig. 6D). These data demonstrate that NbSYN87 is only selective against the human but not the mouse α-synuclein protein.

**Figure 6:**
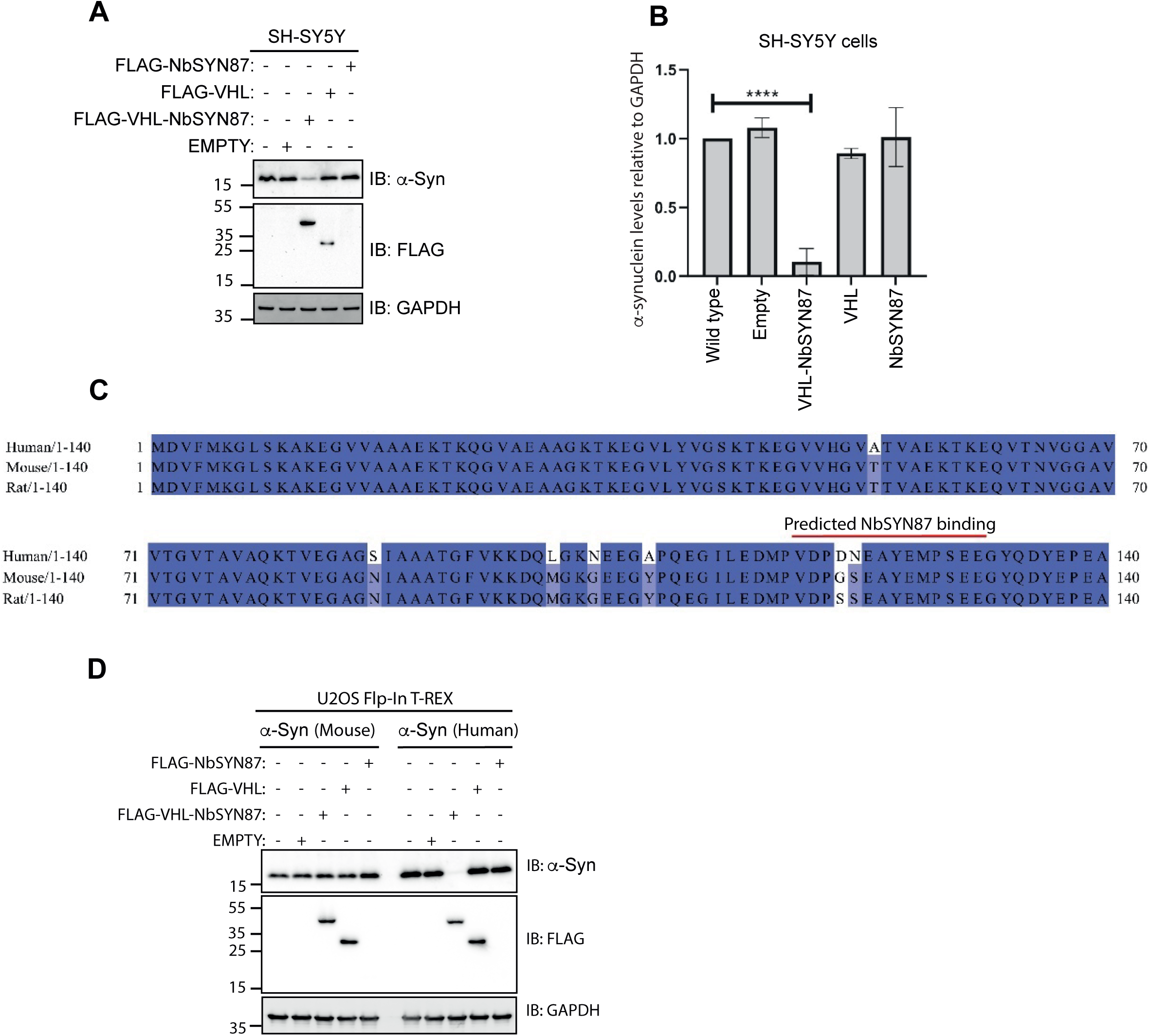
The VHL-NbSYN87 AdPROM degrades human but not mouse α-synuclein. **A)** SH-SY5Y neuroblastoma cells were retrovirally transduced to express either VHL-NbSYN87, VHL, NbSYN87 or empty vector control. 20 μg of extract protein was resolved by SDS-PAGE and transferred onto nitrocellulose membranes, which were subjected to immunoblotting with the indicated antibodies. Blots representative of at least n=3 independent experiments. **B)** Quantification of α-synuclein levels from (A) relative to loading GAPDH control ±SD of n=3 independent experiments. **C)** Alignment of human, mouse and rat α-synuclein sequences carried out using Jalview software. Likely epitope of NbSYN87 is indicated by the red line. **D)** U2OS Flp-In T-Rex cells expressing either mouse or human α-synuclein were retrovirally transduced to express FLAG-tagged VHL, NbSYN87, VHL-NbSYN87 or empty vector control. Cell extracts (20 µg) were resolved by SDS-PAGE and transferred to nitrocellulose membranes for immunoblotting with the indicated antibodies. Blots representative of n=3 independent experiments.

Next, we sought to assess the extent and consequences of VHL-NbSYN87-mediated degradation of α-synuclein on physiologically relevant primary cortical neurons derived from rodents. Using lentiviral vectors, these neurons can be transduced to express exogenous transgenes. Furthermore, these neurons can also be seeded with recombinant human α-synuclein pre-formed fibrils (PFFs) to induce aggregation of α-synuclein^35^. Having established that NbSYN87 does not recognise rodent α-synuclein, cortical neurons were transduced with lentiviral vectors encoding human α-synuclein to assess VHL-NbSYN87-mediated degradation of exogenously expressed human α-synuclein in this neuronal model.

### Targeted degradation of α-synuclein delays PFF-induced aggregation in primary cortical neurons

Rat cortical neurons were co-transduced with varying dilutions of lentiviral vectors encoding the expression of NbSYN87 (starting at MOI 13.3), VHL (starting at MOI 10.4), VHL-NbSYN87 (starting at MOI 8.65) or empty control (starting at MOI 12.5) with all also encoding non-fused mKate2 transgene used as an expression control together with a lentiviral vector encoding the expression of human WT α-synuclein (MOI 0.33). There was no interference with the mKate2 fluorescence under any experimental condition (Fig. 7A). Empty mKate2 and no treatment controls served as negative controls. Primary cortical neurons overexpressing human WT α-synuclein displayed substantially reduced levels of human WT α-synuclein signal with the VHL-NbSYN87 AdPROM at DIV9 in comparison to NbSYN87, VHL, empty mKate2 and no treatment controls (Fig. 7B). In line with these results, biochemical analysis using MSD ELISA demonstrated a significant reduction of total human (Fig. S5A) α-synuclein levels induced by the VHL-NbSYN87 AdPROM under non-seeding conditions when compared to the NbSYN87, VHL and empty mKate2 lentiviral controls transduced at the highest MOIs. No significant changes in levels of rodent α-synuclein were observed under any treatment conditions (Fig. S5B). In primary cortical neurons overexpressing human WT α-synuclein and seeded with recombinant human α-synuclein PFFs (8 ng/ml), co-transduction with the VHL-NbSYN87 lentiviral vector resulted in substantially reduced aggregated pS129-α-synuclein signal at DIV16 relative to NbSYN87, VHL, empty mKate2 and no treatment controls (Fig. 7C), suggesting inhibition of seed-induced aggregation of α-synuclein in cortical neurons by the VHL-NbSYN87 AdPROM. These data were confirmed using biochemical MSD ELISA analysis showing significantly reduced aggregated phospho-S129 (Fig. 7D) and substantially decreased total human α-synuclein levels (Fig. 7E) upon transduction with the VHL-NbSYN87 lentiviral vector in contrast to the NbSYN87, VHL, empty mKate2 and no treatment conditions at the highest MOIs.

**Figure 7:**
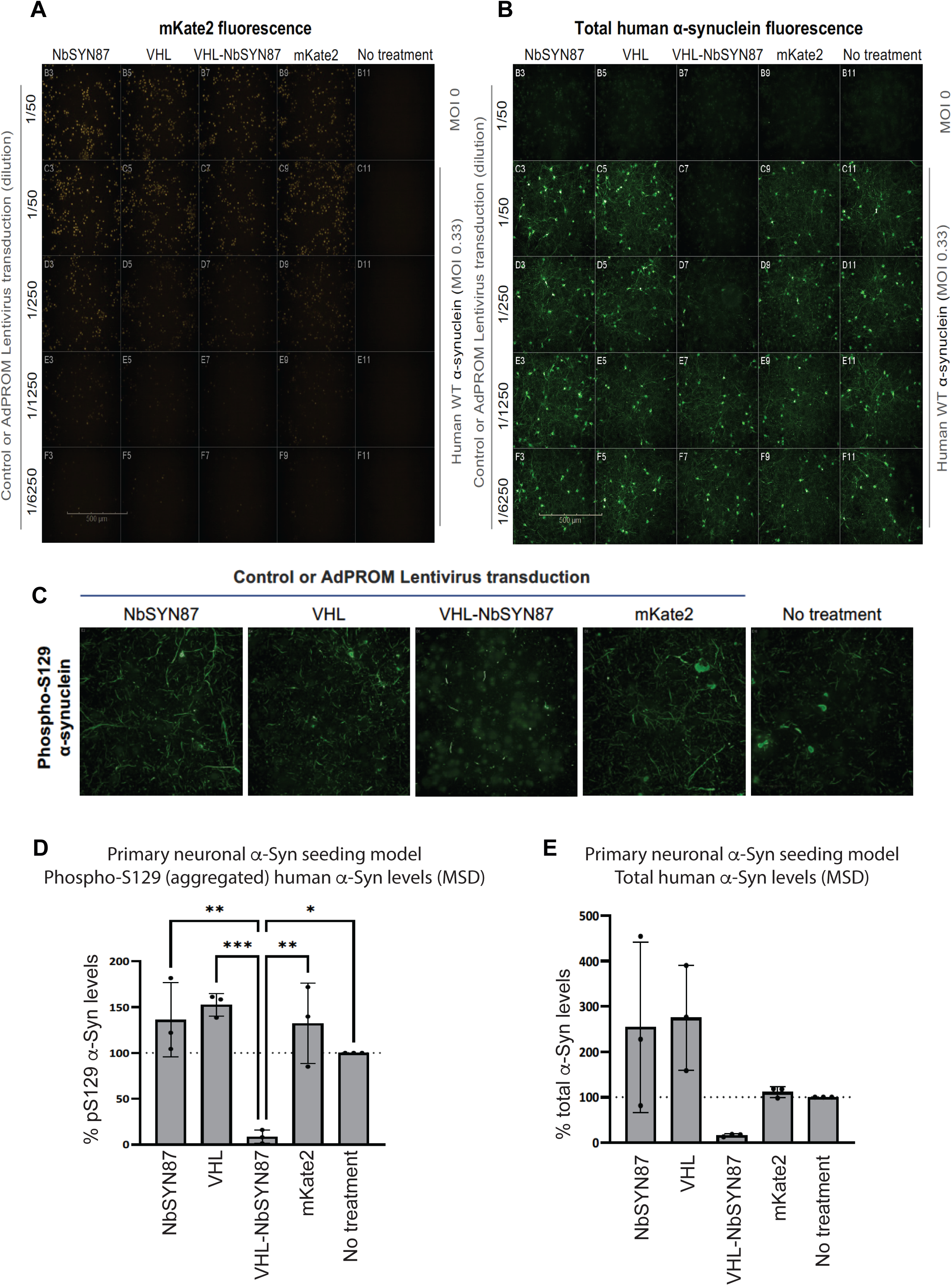
Targeted degradation of α-synuclein delays PFF-induced aggregation in primary cortical neurons. **A)** Representative photomicrographs of mKate2 fluorescence on primary cortical neurons in absence (MOI 0, top panel) or presence of lentiviral vector-mediated overexpression of human WT α-synuclein (MOI 0.33, rest) transduced together with the indicated dilutions of lentiviral vectors encoding NbSYN87, VHL, VHL-NbSYN87, empty vector with non-fused mKate2 (orange) or no treatment condition. Scale bar represents 500 µM. **B)** As in (A) except the primary cortical neurons were subjected to immunofluorescence with an antibody recognising total human α-synuclein (green) at DIV9. Scale bar represents 500 µM. **C)** As in (B) except the primary cortical neurons overexpressing human WT α-synuclein were seeded with recombinant human α-synuclein PFFs (8 ng/ml) and were subjected to immunofluorescence with an antibody recognising phospho-S129 (aggregated) α-synuclein (green) at DIV16. **D&E)** MSD analysis of phospho-S129 (aggregated) α-synuclein **(D)** and total human α-synuclein **(E)** for samples from (C). Results are shown as mean ± SD (3 biological replicates with 2 technical replicates). Statistical analysis was carried out using ordinary one-way ANOVA, Bonferonni’s multiple comparisons test.***, P-value < 0.001; **, P-value < 0.01; *, P-value < 0.05. αSyn: α-synuclein

## Discussion

In this manuscript, we report a VHL-NbSYN87 AdPROM that efficiently and selectively targets the degradation of human α-synuclein from cancer-derived cell lines and primary rat cortical neurons expressing human α-synuclein. Furthermore, the expression of the VHL-NbSYN87 AdPROM prevented the accumulation of α-synuclein aggregates in rat cortical neurons when seeded with recombinant human α-synuclein PFFs. The AdPROM-mediated inhibition of PFF-induced accumulation of toxic α-synuclein aggregates that we show here demonstrates the potential for targeted α-synuclein degradation to reverse toxicity associated with aggregation. Previous studies carried out in dopaminergic neurons differentiated from iPSCs from a *SNCA* triplication patient showed that genetic ablation of α-synuclein reduced seeded aggregation, compared to *SNCA*-triplicated control cells^23^. Other studies using *SNCA*-KO cells have shown that ablation of α-synuclein protein levels results in resistance to certain neurotoxic models of Parkinsonism^36–38^. Similarly, RNAi-mediated depletion of *SNCA* transcripts have shown promise in mitigating disease progression^39^. Indeed, BIIB101, a *SNCA*-targeting anti-sense oligonucleotide developed by Ionis and Biogen is currently undergoing a Phase I clinical trial in MSA patients (NCT04165486)^40^. These studies, along with our observations here, suggest that a reduction in α-synuclein protein levels may be able to halt or slow disease progression in synucleinopathies. Therefore, targeted degradation of α-synuclein could offer a disease-modifying therapeutic opportunity for synucleinopathies.

Targeted degradation has emerged as an exciting modality to target the destruction of disease-causing proteins^26,27^. Several PROTACs have entered clinical trials and many molecular glue degraders are already used in clinical settings^27^. In the absence of potent and selective small molecule binders of α-synuclein, currently no PROTACs or molecular glues are available for targeting the degradation of α-synuclein. Nonetheless, our work demonstrates that targeted degradation of α-synuclein through the proteasome can be achieved and is also effective at preventing seed-induced aggregation of α-synuclein. Moreover, the VHL-NbSYN87 AdPROM is capable of degrading all clinically relevant mutants of α-synuclein. The next leap would be to test whether the VHL-NbSYN87 is also effective at clearing pre-formed aggregates from affected brains in *in vivo* models of synucleinopathies. If effective, delivering the VHL-SYN87 AdPROM to the affected neurons through AAVs could be used for therapeutic targeting of α-synuclein aggregates. Encouragingly, when two α-synuclein nanobodies, including NbSYN87, were tethered to a proteasome-targeting sequence, this led to a partial clearance of α-synuclein from cells overexpressing α-synuclein^41^ and when inoculated into the brain of an α-synuclein-based PD rodent model, pathologic aggregation of α-synuclein was shown to be reduced^42^. We would anticipate much better responses in similar assays by the VHL-NbSYN87 AdPROM, given the robust degradation of α-synuclein we have demonstrated in the clonal and primary neuronal cell models. Similarly, a recent study reported the development of ATC161, an autophagy targeting chimera that targeted the degradation of α-synuclein aggregates, but not soluble α-synuclein, through the autophagy-lysosomal pathway and displayed therapeutic efficacy against synucleinopathy-associated genotoxicity and mitotoxicity^43^.

The VHL-NbSYN87 AdPROM relies on the CUL2-Rbx1 E3 ligase machinery to cause the degradation of α-synuclein via the ubiquitin-proteasome pathway. The AdPROM-mediated degradation of α-synuclein could be further improved by replacing VHL with any E3 ubiquitin ligase that displays activity within dopaminergic neurons and by replacing NbSYN87 with nanobodies or intrabodies that selectively bind to toxic, pathogenic mutant, or aggregated forms of α-synuclein. Despite the long time that has passed since mutations and aggregation of α-synuclein in the pathogenesis of PD and other synucleinopathies have been established, therapies targeting α-synuclein have been slow to progress^6^. However, targeted degradation of α-synuclein offers much needed hope for progress in therapeutic targeting of α-synuclein.

## Materials and Methods

### Materials

The siRNA oligonucleotides used were purchased from Dharmacon. The product reference for the siRNA targeting human α-synuclein (SNCA) is L-011109-00-0005. The non-targeting control siRNA reference is D-001810-01-05.

### Plasmids

All DNA constructs were generated by MRC PPU Reagents and Services and sequenced by MRC PPU DNA Sequencing and Services. All plasmids used in this study are available upon request at https://mrcppureagents.dundee.ac.uk. For SK-MEL-13 *SNCA*-KO cells, the following guide RNAs (gRNAs) were used: sense gRNA (DU64505) and antisense gRNA (DU64516).

### Cell culture

U2OS osteosarcoma, HeLa, human embryonic kidney HEK 293, retroviral production HEK293 FT, A549 lung adenocarcinoma, SK-MEL-13, SK-MEL-23 and G-361 melanoma cells were grown in DMEM supplemented with 10% (v/v) FBS, 2 mM L-glutamine and 100 U/ml pen/strep (referred to as full medium hereafter). SH-SY5Y neuroblastoma cells were cultured in DMEM/F-12 supplemented with 10% (v/v) FBS, 2 mM L-glutamine and 100 U/ml pen/strep.

Cells were maintained at 37°C in a humidified incubator set to 5% CO_2_. Aseptic conditions were used when performing all cell culture procedures. Cells were grown to 80-90% confluency in a 10- or 15-cm Nunclon-coated dish (Thermo Fisher Scientific). For passaging, cell medium was aspirated from the dish and the cells were washed once in PBS. The PBS was then aspirated from the dish and the cells were incubated with 1-2ml of trypsin/EDTA at 37°C until all cells had detached from the dish. Detached cells were resuspended in fresh full medium and pipetted a few times to detach cell clumps before being seeded into new culture dishes at the desired dilutions/densities. For cell counting, a CellDrop automated cell counter (DeNovix) was used as per manufacturer’s instructions. Extracts from other cell lines that were used to analyse α-synuclein expression were obtained from various labs within the MRC-PPU. When using RNAi technology to knockdown protein levels, cells were seeded at 2×10^5^ cells in a 6-well culture dish 24 hours before transfection. To transfect, 150 µl Opti-MEM and 5 µl of Lipofectamine RNAiMAX were mixed in one tube whilst 150 µl Opti-MEM and 1 µl of siRNA (made to a 20 µM stock so 1 µl is equal to 20 pmol) were mixed. The two tubes were mixed together gently and left to incubate for 20 minutes at room temperature. The solution was added dropwise to the wells and left for 24 hours. After 24 hours, media was changed and the cells were left for a further 24 hours before lysis.

### Retroviral generation of stable cell lines

Retroviral vector was generated using pBABED-puro vectors as described previously^44^. 6 μg of vector was transiently transfected into a 10-cm dish of approximately 70% confluent human embryonic kidney (HEK) 293FT cells alongside 2.2 μg of pCMV5-VSV-G and 3.8 μg of pCMV5-GAG/POL. Briefly, the plasmids were added to 300 μl of Opti-MEM in one tube. 24 μl of 1 mg/ml PEI was added to 300 μl of Opti-MEM in a second tube. Both tubes were left to incubate for 5 minutes before being mixed together and left for a further 20 minutes. Contents of the tube were then added to 9 ml of complete DMEM media before being placed onto the HEK 293FT cells. The transfection medium was replaced with fresh medium after 16 hours. A further 24 hours later, the viral media was collected and passed through a 0.45 μm sterile syringe filter. Target cells at ∼60% confluency were transduced with the optimised titre of the retroviral medium diluted in fresh medium containing 8 μg/ml polybrene (Sigma-Aldrich) for 24 hours. The retroviral medium was then replaced with selection medium containing 2 μg/ml puromycin for selection of cells transduced with the intended retroviral particles. A pool of transduced cells were utilised for subsequent experiments following complete death of non-transduced cells placed under the selection medium in parallel.

### Generation of Flp-In T-Rex cell lines

Flp-In T-Rex U2OS and HeLa cells were maintained in complete medium supplemented with 15 μg/ml blasticidin and 100 μg/ml zeocin to maintain expression of the Tet-repressor and integrity of the Flp-recombination site respectively. Cells at ∼60-70% confluency were transfected with 1 μg of the pcDNA5-FRT/TO vector encoding for the POI as well as 9 μg of the pOG44 Flp recombinase plasmid in 1 ml of Opti-MEM with 20 μl of 1 mg/ml PEI . The transfection mixture was incubated at room temperature for 20 minutes before being added dropwise to the cells. After 24 hours, media was replaced with fresh selection media containing 15 μg/ml blasticidin and 50 μg/ml hygromycin-B. The selection medium was replaced every 2-3 days for approximately 2-3 weeks until resistant clones were selected. Clones were expanded and verified by immunoblotting. To induce protein expression, cells were incubated with 20 ng/ml doxycycline for 24 hours prior to use, unless stated otherwise in figure legends. In some cases, leaky levels of expression were seen and these were used to reduce any artefacts potentially caused by overexpression of the POI.

### Generation of SK-MEL-13 SNCA-KO cell lines using CRISPR/Cas9 technology

CRISPR/Cas9 gene editing technology was used to generate *SNCA*-KO SK-MEL-13 cells. Briefly, SK-MEL-13 cells were transfected at approximately 60-70% confluency. In 10-cm culture dishes, 1 ml of Opti-MEM and 20 µl PEI were combined with 1 µg of each of the SNCA-targeting sense or antisense guide-RNA plasmids, which also encoded Cas9-D10A and puromycin-resistance, respectively. This mixture was vortexed for 20 seconds and left standing at room temperature for 20 minutes before being added dropwise to the cells. Cells were left for 24 hours before medium was changed to selection medium containing 2 µg/ml puromycin. Selection medium was also added to a plate of control cells that had not been transfected as a negative control. Cells were left in selection medium for 48 hours before being replaced to allow for recovery. The transfection process was repeated but cells were not selected with puromycin. Cells were sorted by fluorescence-activated cell sorting (FACS) to obtain single cell colonies. Cells were prepared for sorting by trypsinisation and resuspension in full medium before being pelleted and resuspended in DMEM with 1% (v/v) FBS. FACS was used to sort the cells into individual wells in 96-well plates which had been pre-coated with 1% (w/v) gelatin and contain 200 µl of conditioned media supplemented with 20% (v/v) FBS. The plate was then briefly centrifuged at 500 xg for 1 minute to ensure no cells got trapped in the meniscus. Viable clones were expanded and *SNCA*-KO clones were analysed and verified by immunoblotting and DNA sequencing.

### Cell lysis

Cells were first washed twice in ice-cold PBS before being scraped on ice in lysis buffer (50 mM Tris-HCl pH 7.5, 0.27 M sucrose, 150 mM NaCl, 1 mM EGTA, 1 mM EDTA, 1 mM sodium orthovanadate, 10 mM sodium β-glycerophosphate, 50 mM sodium fluoride, 5 mM sodium pyrophosphate and 1% NP-40) supplemented with 1x cOmplete™ protease inhibitor cocktail (Roche). Extract was transferred to Eppendorf tubes and incubated by rotating for 30 minutes to 1 hour at 4°C. Extracts were cleared by centrifugation at 17,000 rpm for 20 minutes at 4°C. Supernatant was transferred to a fresh Eppendorf tube and pellet was discarded. Samples were either processed for use immediately or else snap-frozen in liquid nitrogen and stored at −80°C for later use. For DNA or mRNA isolation, cells were lysed and processed using a Qiagen DNeasy blood and tissue kit or RNeasy kit respectively, as per the manufacturer’s guidelines.

### SDS-PAGE and Western Blotting

Cell extracts containing equal amount of protein (typically 10-20 μg as indicated) were resolved by SDS-PAGE using Bis-tris gels. Samples were transferred to nitrocellulose membranes. For immunoblotting, membranes were blocked for 1 hour at room temperature with 5% (w/v) non-fat milk (Marvel) in TBS-T (20 mM Tris, 150 mM NaCl, 0.1% Tween-20). Membranes were incubated overnight at 4°C in 5% (w/v) milk/TBS-T containing a dilution of the appropriate primary antibody. Primary antibodies and the dilutions they were used at are as follows: anti-α-synuclein (Ab6162, Abcam, 1:500), anti-FLAG HRP (A8592, Sigma, 1:1000), anti-HIF1α (610959, BD Biosciences, 1:1000), anti-GAPDH (2118S, CST, 1:5000) anti-GFP (11814460001, Sigma, 1:1000), anti-Cullin2 (51-1800, Invitrogen, 1:1000), and anti-ubiquitin (Z0458, DAKO, 1:1000). Membranes were subsequently washed with TBS-T and incubated for 1 hour at room temperature with HRP-conjugated or fluorescent secondary antibodies. Secondary antibodies used and their dilutions are as follows: rabbit anti-sheep-IgG (31480, Thermo Fisher Scientific, 1:2500), goat anti-rabbit-IgG (7074, CST, 1:5000), goat anti-mouse-IgG (31430, Thermo Fisher Scientific, 1:5000), StarBright Blue 700 goat anti-rabbit-IgG (12004161, Bio-Rad, 1:5000). Membranes were again washed in TBS-T and signal detection was performed using ECL (Merck) and the ChemiDoc MP System (Bio-Rad). Image Lab (Bio-Rad) was used to analyse protein bands by densitometry.

### Immunofluorescence microscopy (for SK-MEL-13 cells)

For immunofluorescence imaging, cells were first seeded onto 16-mm diameter circular sterile coverslips in 12-well culture plates and left to adhere overnight. All coverslips were sterilised with 100% (v/v) ethanol and allowed to dry prior to use. Cells were washed twice in PBS before being fixed in 4% (w/v) paraformaldehyde (diluted in PBS) at room temperature for 10 minutes.

PFA was removed and the coverslips were washed twice in PBS. Coverslips were then permeabilised with 0.2% (v/v) NP40 in PBS for 3 minutes. Cells were then blocked by washing twice and a 15-minute incubation in 1% (w/v) BSA/PBS. Coverslips were then incubated for 1.5 hours at room temperature with a 1:100 dilution of anti-α-synuclein antibody (610786, BD Biosciences) in a humidified chamber. Coverslips were then washed 3 times (for 10 minutes each) in 0.2% (w/v) BSA/PBS before being incubated with goat anti-mouse-IgG Alexa Fluor 594 conjugated secondary antibody (A-11005, Thermo Fisher Scientific) at a dilution of 1:500 for 1 hour at 37°C, protected from light. Coverslips were then washed three times in 0.2% (w/v) BSA/PBS for 10 minutes each with the first wash containing a 1:15,000 dilution of DAPI. They were then briefly immersed in dionised water using a tweezers and placed on a paper towel to air dry. Once dry, approximately 5 μl of Vectashield was dotted onto a glass slide and the coverslip was gently laid (cell side down) onto the solution before being sealed with clear nail polish. Cells were imaged on a Deltavision system (Applied Precision) with an immerse-oil 60x or 40x objective and processed with SoftWoRx (Applied Precision). Where applicable, Z-series were obtained and deconvolved using SoftWoRx. Images were processed and figures were made using Adobe Photoshop or OMERO software.

### Proteomic analysis

#### Cell Lysis, in-solution digestion, TMT labelling and fractionation

The cells were lysed in lysis buffer (8 M urea, 20 mM HEPES pH 8.0,1 mM sodium orthovanadate 2.5 mM sodium pyrophosphate,1 mM ß-glycerophosphate), sonicated and centrifuged at 16,000 xg for 20 minutes. Protein concentration was determined using the BCA assay (Pierce, Waltham, MA). From each sample, 100 µg of protein was reduced with 5 mM DTT for 20 minutes at RT and alkylated with 10 mM iodoacetamide for 10 minutes at room temperature in the dark. For trypsin digestion, samples were diluted to reduce the urea concentration to 2 M with 20 mM HEPES, pH 8.0 and subjected to digestion with TPCK-treated trypsin in 1:20 enzyme to substrate ratio (Worthington Biochemical Corp, Lakewood, NJ) for 12–16 hours at RT. Digested peptides were acidified by 1 % trifluoroacetic acid (TFA) and desalted using C18 Sep-Pak cartridge (Waters, Cat#WAT051910) and dried in SpeedVac vacuum concentrator. These peptides from corresponding samples were labelled with 16-plex TMT reagents (Thermo Fisher) according to the manufacturer’s instructions. The labelled peptides were pooled together and subjected to basic pH fractionation. Briefly, peptides were dissolved in Buffer A (10 mM ammonium formate, pH 10) and resolved on an XBridge BEH RPLC column (Waters XBridge BEH C18 Column, 130 Å, 5 µm, 4.6 mm × 250 mm. #186003010) at a flow rate of 0.3 ml/minute by applying gradient of 7–40% (solvent-B, 90% by vol. acetonitrile in 10 mM ammonium formate, pH 10) for 80 minutes into a total of 96 fractions. Each adjacent fraction was pooled together to make 48 fractions for mass spectrometry analysis and dried on SpeedVac vacuum concentrator.

### Mass spectrometry analysis

Individual peptide fractions were reconstituted in 0.1% formic acid and analysed on an Orbitrap Fusion Tribrid Mass spectrometer (Thermo Fisher Scientific, San Jose, U.S.A.) interfaced with the Dionex 3000 RSLC nano-liquid chromatography. The peptide samples were enriched on a nano viper trap column (C18, 5 µm, 100 Å, 100 µm × 2 cm; PN: 164562, Thermo Fisher Scientific) and separated on a 50 cm analytical column (2 µm, 100 Å, 75 µm × 50 cm; PN: ES803, Thermo Fisher Scientific) by the gradient of Solvent B (100% CAN, O.1% formic acid) for 140 minutes. The mass spectrometer was operated in a Data-Dependent Acquisition (DDA) mode using SPS MS3 method. A survey full scan MS (from m/z 375-1500) was acquired in the Orbitrap with a resolution of 120,000 at 200 m/z. Top speeds comprising 3-second cycle time were used. The AGC target for MS1 was set as 4×10^5^ and ion filling time was set at 50 ms. The precursor ions for MS2 were isolated using a Quadrupole mass filter at a 0.7 Da isolation width, fragmented using a normalized 35% HCD of ion routing multipole and analysed using ion trap. The top 10 MS2 fragment ions in a subsequent scan were isolated and fragmented using HCD at 65% normalized collision energy and analysed using an Orbitrap mass analyser with a resolution of 50,000 at scan range of 100-500 m/z. Dynamic exclusion was set for 30 seconds with a 10 ppm mass window.

### Data analysis

Raw MS data were searched using the MaxQuant search algorithm (version 2.0.3.0) against the Uniprot human protein database. A 10-plex TMT reporter ion MS3 workflow was loaded and used following the search parameters. Trypsin as a protease was selected by allowing two missed cleavages, deamidation of Asn and Gln, oxidation of Met was used as a variable modification, and carbamidomethylation of Cys was used as a fixed modification. The default mass error tolerance for MS1 and MS2 (4 ppm and 20 ppm) was used. A minimum of two unique + razor peptides were selected for quantification. The data were filtered for 1% PSM and peptide and protein level FDR. For the identification of significantly differential proteins, two-sample statistical “t-test” was performed.

### Primary neuronal α-synuclein assays: Primary rat cortical neuronal cultures

Rat primary cortical neurons were isolated from the cortex of Wistar rat embryos (Janvier) following ethical guidelines. All experimentation at Johnson & Johnson involving laboratory animals was approved by the Institutional Ethical Committee on Animal Experimentation in accordance with EU-directive 2010/63/EU and the Belgian Royal Decree of 29 May 2013. The facility was and is AAALAC accredited. Neurons were plated in 96-well plates (655946, Greiner Bio-One) previously coated with poly-L-lysine (P1274, Sigma-Aldrich) at a density of 40,000 cells per well. Cells were cultured at 37°C and 5% CO_2_ in Neurobasal medium (10888022, Gibco) supplemented with 5% B-27 (17504044, Invitrogen) and 2.5% GlutaMax (35050038, Invitrogen). At DIV0, cells were transduced with human WT α-synuclein lentiviral vector (MOI 0.33 under non-seeding conditions; MOI 1 under seeding conditions) under the control of the human synapsin 1 promoter (VectorBuilder). At DIV2, cells were treated with recombinant human α-synuclein pre-formed fibrils (PFFs; 8 ng/µl, Johnson & Johnson Innovative Medicine) and/or transduced with lentiviral vectors encoding NbSYN87 (starting at MOI 13.3), VHL (starting at MOI 10.4), VHL-NbSYN87 (starting at MOI 8.65) or empty control which also encoded non-fused mKate2 (starting at MOI 12.5) under the control of the human synapsin 1 promoter (VectorBuilder). Cells were fixed in 4% paraformaldehyde for immunocytochemical analysis or lysed in RIPA lysis buffer for biochemical analysis at DIV9 (non-seeding conditions) or DIV16 (seeding conditions).

### Immunocytochemical staining

Fixed cells were blocked in PBS-0.1% Triton-X100 with 10% donkey serum (D9663, Sigma-Aldrich) or 5% normal goat serum (G9023, Sigma-Aldrich) for 1 hour. Cells were then incubated overnight in PBS-0.1% Triton-X100 with 10% donkey serum or 5% normal goat serum and primary antibodies against rabbit α-synuclein MJFR1 (Ab209420, 1:2000, Abcam) and mouse phosphorylated (pS129) α-synuclein clone 81A (Ab184674, 1:2000, Abcam). Secondary antibodies used were donkey anti-rabbit Alexa Fluor 488 (A21206, 1:1000, Invitrogen), goat anti-rabbit Alexa Fluor 680 (A21109, 1:1000, Invitrogen) and goat anti-mouse Alexa Fluor 488 (A28175, 1:1000, Invitrogen). Nuclei were stained using Hoechst 33342 (H3570; 1:10000, Invitrogen) diluted in PBS. Immunocytochemical stainings were visualized using Opera Phenix (40x, water objective, Perkin Elmer) and processed using Harmony software.

### Biochemical analysis using Meso Scale Discovery (MSD) ELISA

Coating antibodies 4B12 (807803, 2 µg/ml, Biolegend), rodent α-synuclein (2 µg/ml, Johnson & Johnson Innovative Medicine) and MJFR13 (Ab168381, 0.5 µg/ml, Abcam) were diluted in PBS and dispensed into MSD plates (30 μl per well; L15XA, Meso Scale Discovery) and were incubated overnight at 4°C. After washing with 5 × 200 μl of PBS-0.5%Tween-20, plates were blocked with 0.1% casein in PBS and washed again with 5 × 200 μl of PBS-0.5%Tween-20. After adding samples and standards (both diluted in RIPA buffer), plates were incubated overnight at 4°C under gentle agitation. Plates were next washed with 5 × 200 μl of PBS-0.5%Tween-20, and detection antibodies MJFR1 (Ab209420, 0.5 µg/ml, Abcam), biotinylated D37A6 (74184, 0.5 µg/ml, Cell Signaling Technologies) and 92B (CABT-B1398, 1 mg/ml, Creative Diagnostics) in PBS were added and incubated for 2 hours at room temperature while shaking at 600 rpm. Plates were washed with 5 × 200 μl of PBS-0.5%Tween-20, and SULFO-TAG^TM^ conjugated goat anti-rabbit (R32AB-1, 0.5 µg/ml, Meso Scale Discovery), streptavidin (R32AD-1, 0.1 mg/ml, Meso Scale Discovery) and goat anti-mouse (R32AC-1, 1 µg/ml, Meso Scale Discovery) antibodies in PBS were added and incubated for 1 hour at room temperature while shaking at 600 rpm. After a final wash with 5 × 200 μl of PBS-0.5%Tween-20, 100 μl of 2 X MSD read buffer T (R92TC, Meso Scale Discovery) was added, and plates were read with an MSD reader (MSD SECTOR Imager 6000, Meso Scale Discovery, Gaithersburg, MD). Protein level ratios were calculated by dividing the measurement of a specific sample by the measurement of the high control (no treatment condition).

### Resource Availability

All mass spectrometry data acquired for this study is deposited in the PRIDE database with the accession number PXD060157.

### Raw Data and Code Availability

The data that support the findings of this study are available from the corresponding author upon reasonable request.

### Lead Contact

Further information and requests for resources and reagents should be directed to and will be fulfilled by the Lead Contact, Gopal Sapkota.

## Author Contributions

BC, GG and GS designed and performed experiments, collected and analysed data and contributed to the writing of the manuscript. NK and SR performed some optimisation experiments. TJM designed the strategies for and generated the CRISPR/Cas9 constructs used in this study. JVE performed some experiments and collected data. LDM, AB and DM conceived the project, overviewed the studies and provided essential feedback to the manuscript. GPS conceived and supervised the project, analysed data, and contributed to the writing of the manuscript.

## Acknowledgements

GPS is supported by the UKRI Medical Research Council (grant MC_UU_00018/6 and MC_UU_00038/6), VLAIO PRiND (HBC.2019.2939; awarded jointly to GPS and Johnson & Johnson Innovative Medicine by the Government of Belgium) and the pharmaceutical companies supporting the Division of Signal Transduction Therapy (Boehringer Ingelheim, GSK, Merck-Serono). BC was supported by The Cunningham Trust PhD studentship and MRC PPU. GG, GS, SR and JVE were supported by VLAIO PRiND (HBC.2019.2939). We thank the Sapkota lab members and Neuroscience Discovery teams at Johnson & Johnson for critical appraisal of the data. We thank E. Allen, A. Muir, S. Dalglish, E. Webster and J. Stark for help and assistance with tissue culture, the staff at the DNA Sequencing services (School of Life Sciences, University of Dundee), and the cloning and antibody teams within the MRC-PPU Reagents and Services (University of Dundee, coordinated by J. Hastie). We thank the staff at the flow cytometry facility (School of Life Sciences, University of Dundee) for their invaluable support. We thank the MRC PPU mass spectrometry service team for their help with the project. We thank I. Goris and G. Meulders for generating the primary rat cortical neuron cultures (Neuroscience Discovery, Johnson & Johnson Innovative Medicine). We thank S. De Ren for producing and performing quality control on the recombinant human α-synuclein pre-formed fibrils (Neuroscience Discovery, Johnson & Johnson Innovative Medicine).

## Conflict of Interest Declaration

GG, JVE, LDM, AB and DM are employees and share holders of Johnson & Johnson. SR is a share holder of Astra Zeneca. All other authors have no conflicts to declare.

---

**Figure S1:**
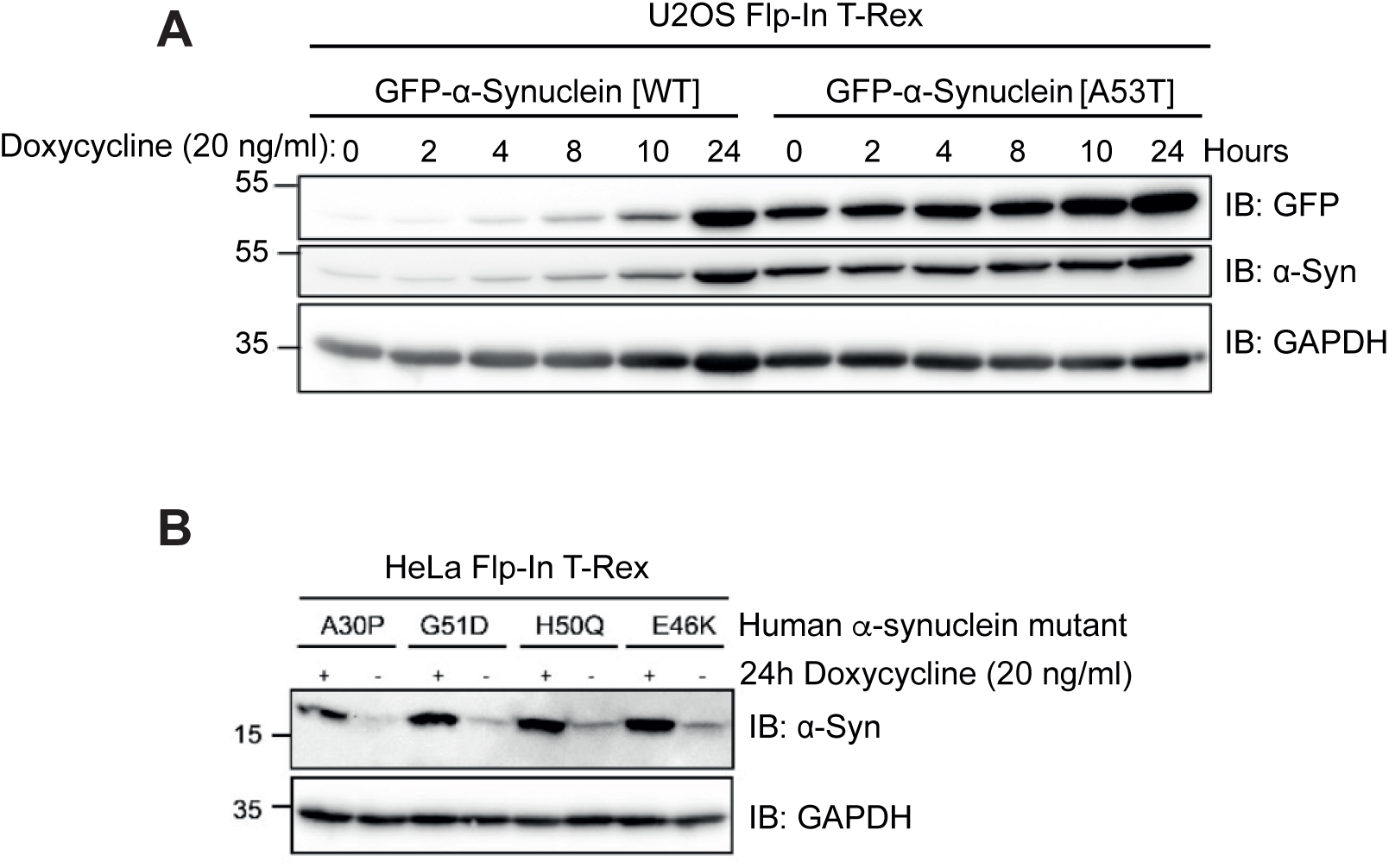
Analysis of Flp IN T-Rex U2OS and HeLa cells expressing WT and mutant α-synuclein. **A)** U2OS Flp-in T-Rex cells stably expressing GFP-tagged WT or A53T mutant human α-synuclein were treated with 20 ng/mL doxycycline for indicated times or left untreated prior to lysis. 20 μg of extract protein was resolved by SDS-PAGE, transferred onto nitrocellulose membranes, which were subjected to immunoblotting with the indicated antibodies. Blots representative of at least n=3. **B)** As in (A) except HeLa Flp-in T-Rex cells stably expressing A30P, H50Q, G51D and E36K mutant forms of utagged human α-synuclein were treated with 20 ng/mL doxycycline for 24 hours or left untreated prior to lysis.

**Figure S2:**
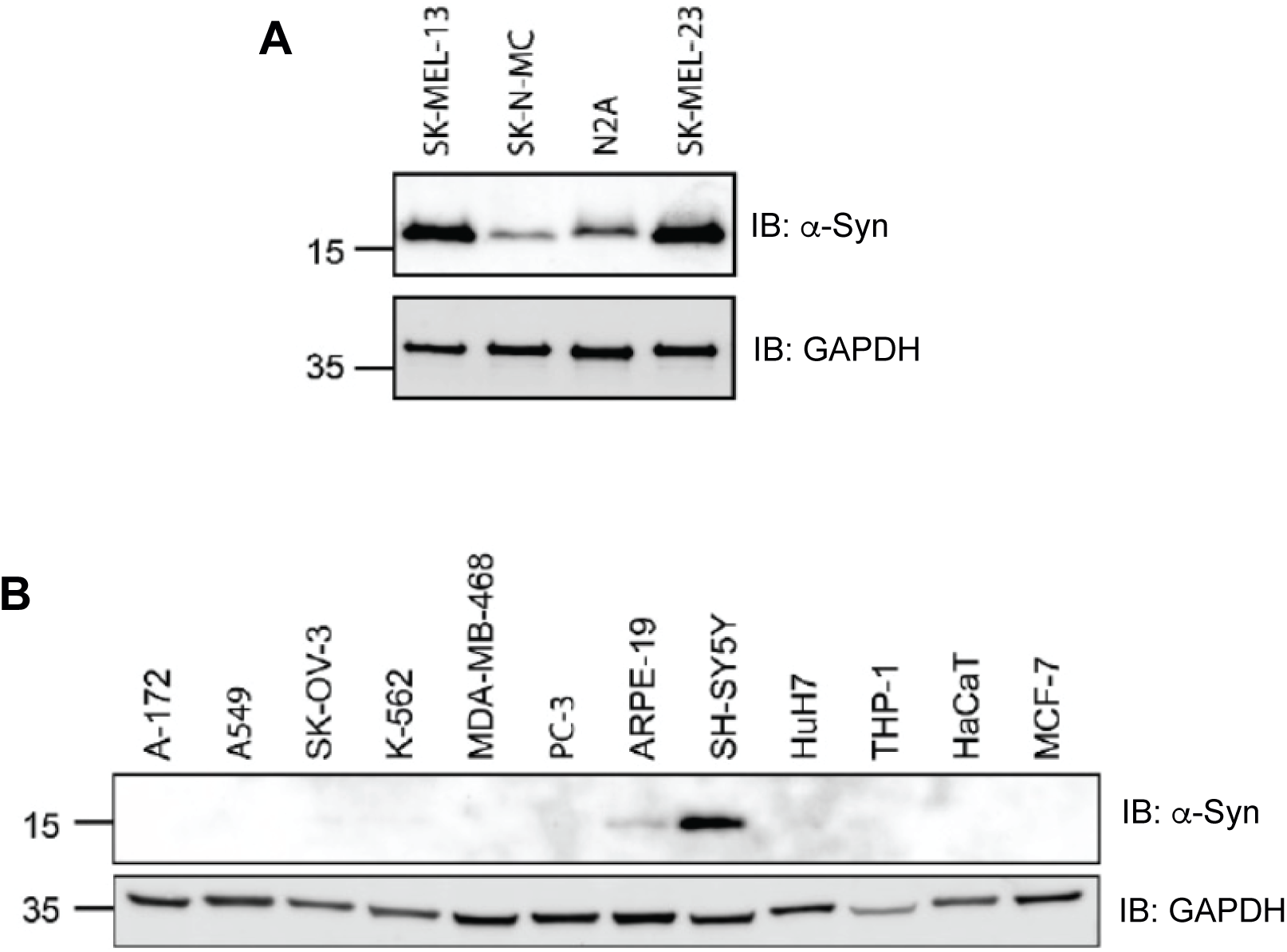
Analysis of α-synuclein abundance in different cell lines. **A)** 20 μg of extract protein each from SK-MEL-13, SK-N-MC neuroblastoma, N2A mouse neuroblast and SK-MEL-23 melanoma cells was resolved by SDS-PAGE, transferred onto nitrocellulose membranes, which were subjected to immunoblotting with the indicated antibodies. **B)** As in (A) except extracts from A-172 glioblastoma, A549 lung adenocarcinoma, SK-OV-3 ovarian cancer, K-562 lymphoblast, MDA-MB-468 breast cancer, PC-3 prostate cancer, ARPE-19 retinal pigment epithelia, SH-SY5Y neuroblastoma, HuH7 hepatoma, THP-1 monocyte, HaCaT keratinocyte and MCF-7 breast cancer cells were used to test α-synuclein abundance.

**Figure S3:**
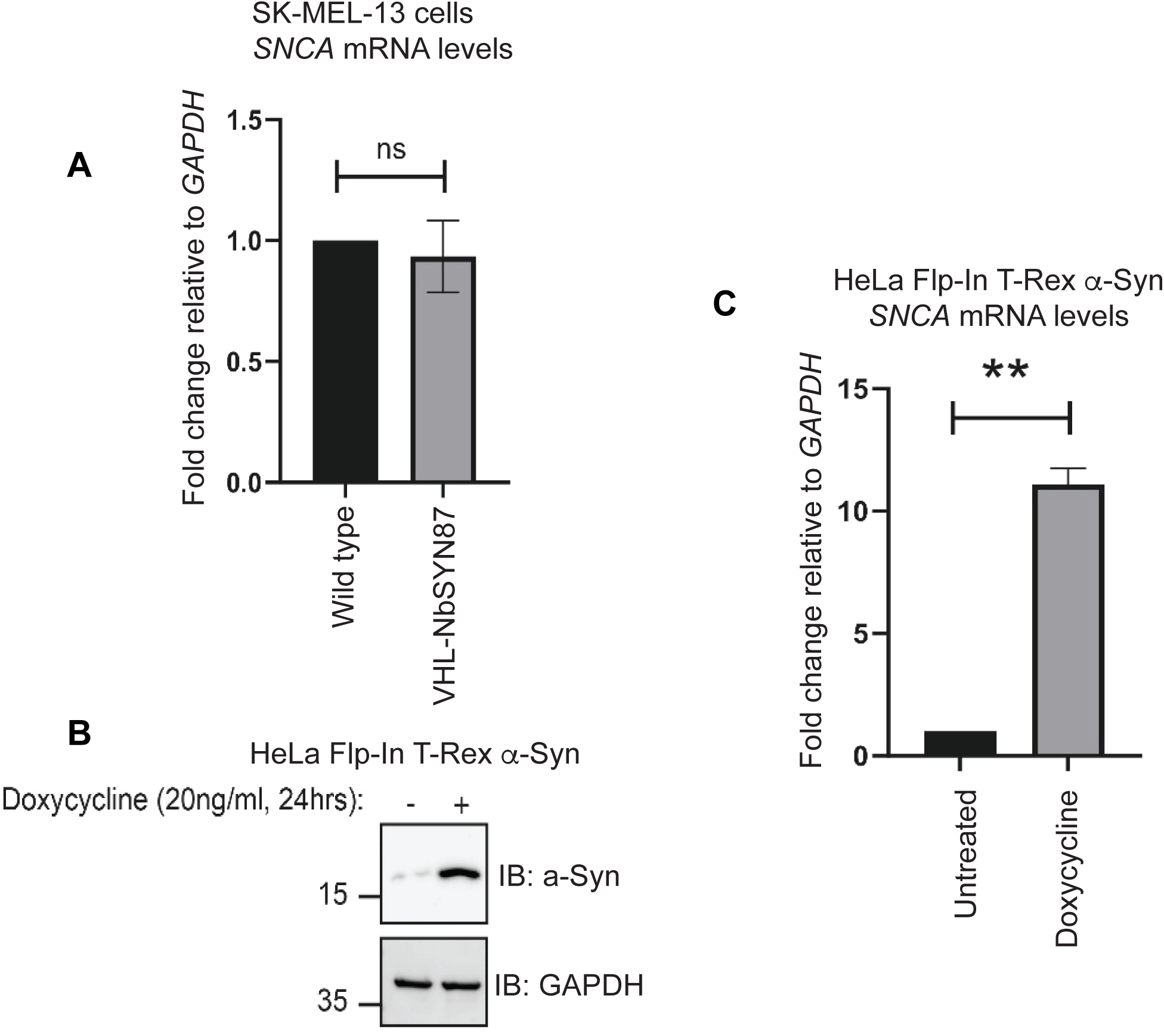
Expression of the VHL-NbSYN87 AdPROM does not affect *SNCA* mRNA levels. **A)** SK-MEL-13 cells retrovirally transduced with empty vector or to express FLAG-VHL-NbSYN87 were subjected to qRT-PCR analysis using primers for *SNCA* and GAPDH. *SNCA* transcript levels relative to internal *GAPDH* levels relative to empty vector control SK-MEL-13 cells are plotted. Error bars show ± SD of n=3 independent experiments. Statistical analysis was carried out using Students T-test. ns, non-significant. **B)** HeLa Flp-In T-Rex α-synuclein cells were treated with or without 20 ng/mL doxycycline for 24 hours before lysis. Cell extracts (20 µg protein) were resolved by SDS-PAGE and transferred to nitrocellulose membranes for immunoblotting with indicated antibodies. **C)** Cells from (B) were subjected to qRT-PCR analysis using primers for *SNCA* and *GAPDH* transcripts to demonstrate that the primers used in (A) do indeed amplify human *SNCA* gene. *SNCA* transcript levels relative to internal *GAPDH* levels relative to control HeLa Flp-In T-Rex cells are plotted. Error bars show ±SD of n=3 independent experiments. Statistical analysis was carried out using Students T-test. **, P value < 0.01.

**Figure S4:**
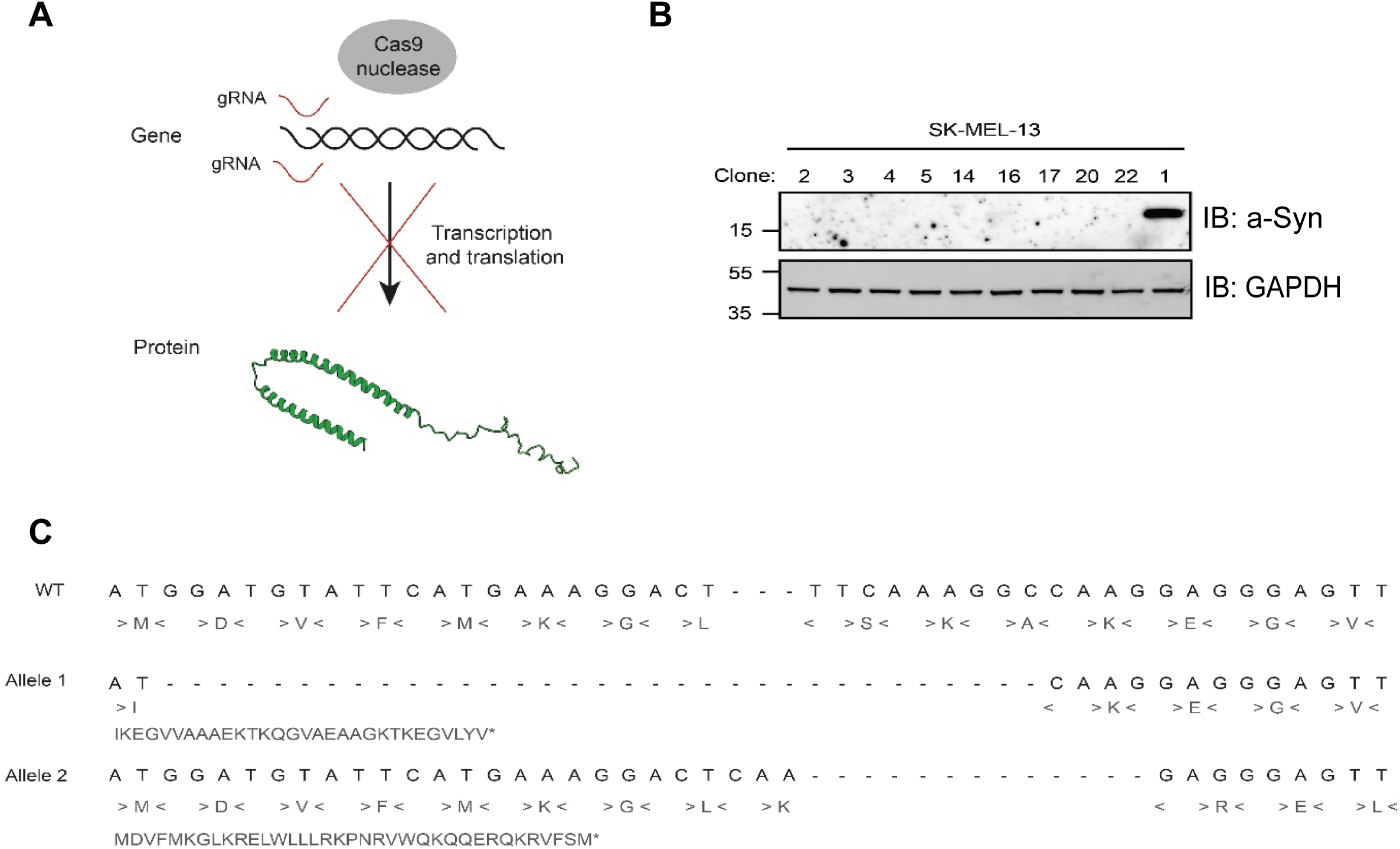
Generation of SK-MEL-13 *SNCA^-/-^* cells using CRISPR/Cas9 genome editing. **A)** Schematic depicting CRISPR/Cas9 methodology. Guide RNAs (gRNAs) are used to guide Cas9 nuclease to the *SNCA* gene to introduce a double-stranded break in the DNA. The repair of this by the cells can create errors that can lead to early stop codons leading to loss of translation of the α-synuclein protein. **B)** A number of single cell clones of potential SK-MEL-13 *SNCA^-/-^* cells were lysed and cell extracts (20 µg protein) were resolved by SDS-PAGE. Proteins were then transferred onto nitrocellulose membranes for immunoblotting with indicated antibodies. **C)** Genomic DNA was isolated from SK-MEL-13 SNCA^-/-^ clone #14 cells with the 5’ end of the SNCA gene being amplified by polymerase chain reaction (PCR) and the DNA being sequenced. The top line shows the DNA sequences, second line shows the corresponding amino acids and the bottom line shows the primary amino acid sequence of the start methionine (M) to the stop codon (*) for both alleles of this KO cell line.

**Figure S5:**
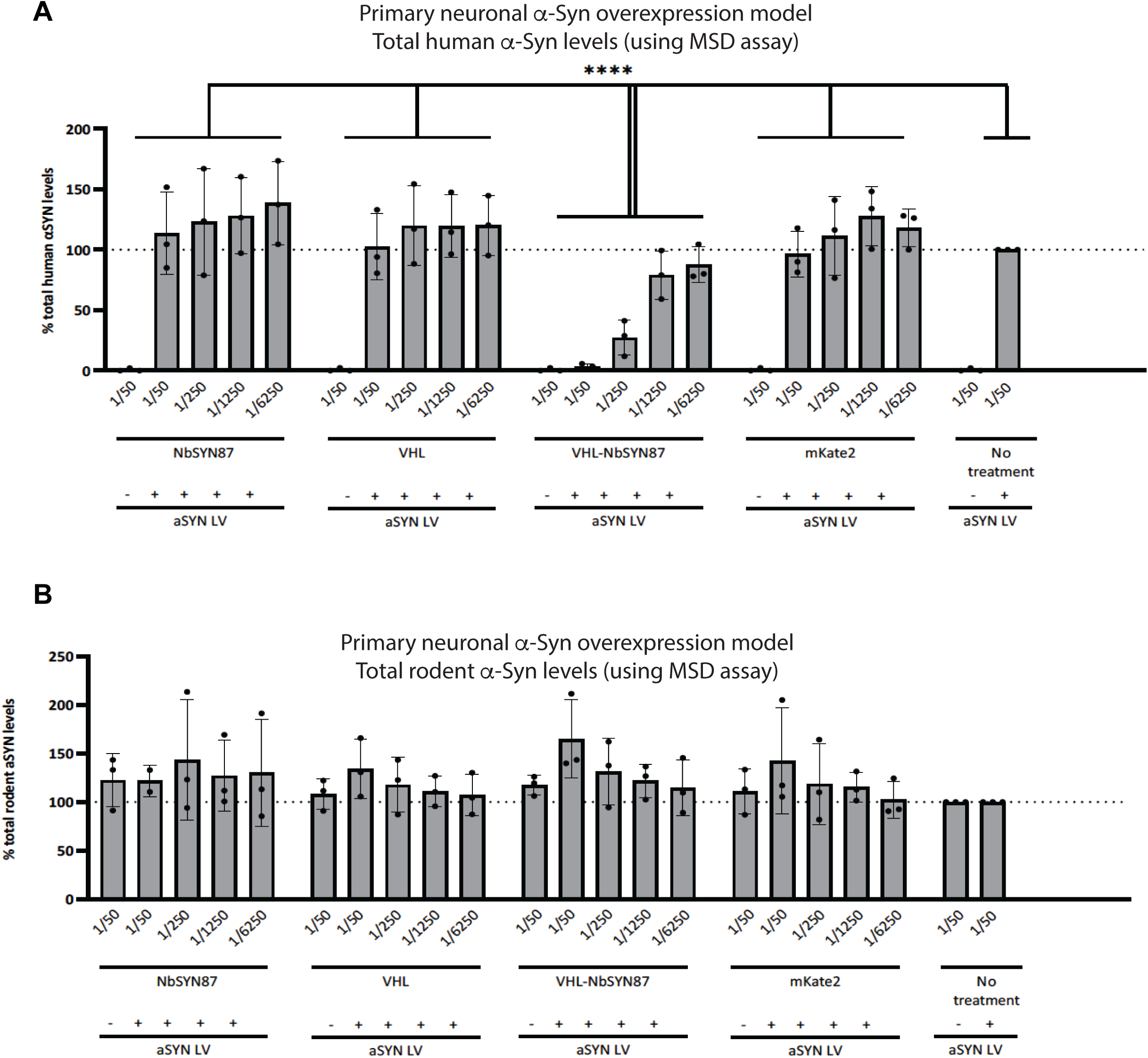
Targeted degradation of only human α-synuclein in primary rat cortical neurons as measured by MSD assay. **A&B)** MSD analysis of total human (A) and rodent (B) α-synuclein levels in primary cortical neurons (from Fig. 7A&B) under non-seeding conditions upon transduction with indicated titres of lentiviral vectors encoding NbSYN87, VHL, VHL-NbSYN87, empty vector with non-fused mKate2 and no treatment condition. Cells were either transduced with lentiviral vectors encoding human α-synuclein (αSYN LV)(+) or not (-) as indicated. Results are shown as mean ±SD (3 biological replicates with 2 technical replicates). Statistical analysis was carried out using ordinary two-way ANOVA, Tukey’s multiple comparisons test. ****, P-value < 0.0001.

